# Physical resilience in the brain: The effect of white matter disease on brain networks in cognitively normal older adults

**DOI:** 10.1101/2022.05.20.492850

**Authors:** Blake R Neyland, Samuel N Lockhart, Robert G Lyday, Laura D Baker, Elizabeth P Handing, Michael E Miller, Stephen B Kritchevsky, Paul J Laurienti, Christina E Hugenschmidt

**Author notes:** Correspondence, **Corresponding Author:** Christina E. Hugenschmidt, Sticht Center on Aging, Wake Forest School of Medicine, 1 Medical Center Blvd, Winston-Salem, NC 27157.

## Abstract

**BACKGROUND:** Physical resilience with age is considered a key feature of healthy aging, but current understanding of the neural contributions to resilience is limited. Additionally, few methods exist to identify physical resilience and observe the mechanisms through which resilience manifests.

**METHODS:** To address these gaps, we used data from 189 participants from the Brain Networks and Mobility (B-NET) study who completed the short physical performance battery (SPPB) as well as its expanded version (eSPPB), magnetic resonance imaging (MRI), and functional MRI (fMRI). Functional brain networks were generated using graph theory methods. We grouped participants based on SPPB scores (<10=unhealthy & 10-12=healthy) and median splits of white matter hyperintensity volumes: Expected Healthy (EH: low WMH, healthy SPPB, n=81), Expected Impaired (EI: high WMH, unhealthy SPPB, n=42), Unexpected Healthy (UH: high WMH, healthy SPPB, n=53), and Unexpected Impaired (UI: low WMH, unhealthy SPPB, n=13). UH is considered the “resilient” group due to their maintained function despite elevated WMH burden. Continuous analyses assessed the relationships between network properties, mobility, and cognition.

**RESULTS:** Higher SPPB scores were associated (p<0.01) with greater sensorimotor cortex community structure (SMN-CS) consistency. While no main effect of the resilience interaction term (SPPB*WMH) was found on SMN-CS, UH showed higher numbers of second-order connections between the SMN and anterior cingulate cortex (ACC) than EI (p<0.01).

**CONCLUSIONS:** Increased connectivity between SMN and ACC may be a marker of physical resilience within the brain.

## INTRODUCTION

The construct of resilience with aging has become an increasingly prominent research focus across many aging related disciplines [1]. The term resilience is commonly defined as the ability to recover from a stressor. Resilience may be considered the opposite of vulnerability and, importantly, is reduced with aging. The majority of research into resilience has focused on cognitive function [2]. Recently, physical resilience, “a characteristic which determines one’s ability to resist or recover from functional decline following health stressor(s)” was proposed as a key feature of healthy aging [1]. A deeper understanding of how physical resilience manifests and allows for the maintenance of physical function may be critical for the promotion of healthy aging in older adults.

With more than 1.5 billion people expected to be over the age of 60 by 2050 [3], age-associated physical decline is expected to significantly affect individual quality of life and financial costs to society [4]. Yet, we do not currently fully understand the development of mobility disability. Recent work suggests that investigating the contributions of the brain and central nervous system to mobility in older adults is important for understanding how mobility disability arises [5], and that aspects of CNS health and organization may contribute to mobility reserve and resilience.

Mobility function requires precise communication between regions across the brain, meaning a disruption to either the connections between regions or within the regions themselves could affect mobility. Structural changes in the brain, such as lower regional gray matter volumes, are associated with mobility impairments in healthy aging adults [6, 7]. Additionally, higher volumes of white matter hyperintensities (WMH) are associated with decreased gait speed [8, 9]. WMH are common magnetic resonance imaging (MRI) findings in older adults [8] and are characterized by damage to the parenchyma, which can result in demyelination or outright axonal disruption [10]. The exact pathogenesis of WMH is not currently known, but they are thought to result from cerebrovascular disease or chronic ischemia [11]. While there is an established relationship between WMH and mobility, it is clear this relationship is neither simple nor linear. For example, there are instances where individuals have high burdens of WMH yet they display no cognitive or physical impairments [12]. This ability to maintain function despite increased WMH accumulation may be indicative of resilience.

One way the brain may adapt to increased WMH volume is through shifts in functional connectivity, i.e., how the different regions of the brain communicate and coordinate to perform tasks. Increased WMH volumes have been associated with both local and distal disruptions in functional connectivity, which may highlight a global network response to pathology [13, 14]. In the context of cognitive reserve and resilience, regional analyses suggest that intrinsic connectivity of the default-mode network (DMN) during rest may play a role in the development of cognitive reserve and resilience with aging [15, 16]. Similarly, changes in functional brain networks have been associated with mobility function in older adults. We have shown that reduced consistency of sensorimotor network community structure (SMN-CS) is associated with lower scores on the short physical performance battery (SPPB) [17], and resting-state functional connectivity has also been associated with walking speed [18]. Research into the neural correlates of physical resilience has been limited, but connectivity of the anterior cingulate cortex (ACC), supplementary motor area (SMA), posterior parietal cortex (PPC), and the cerebellum have recently been implicated in a “fine-tuning” network of mobility [19]. Work to date has not addressed the role of WMH in mobility function or the neural mechanisms of physical resilience.

In an attempt to address the gaps in the literature on the neural correlates of physical resilience, we used graph-theory network methods to assess patterns of functional connectivity that may elucidate compensatory mechanisms indicative of physical resilience. We used baseline data from the Brain Networks and Mobility (B-NET) study, an ongoing observational study on the neural correlates of aging and mobility. To assess reserve and resilience in this cohort, we grouped participants by both WMH volume and scores on the SPPB such that individuals with high WMH volumes and high SPPB scores were deemed resilient. We hypothesized that individuals with high resilience (high SPPB scores while having high WMH burden) would show compensatory shifts in functional connectivity when compared to individuals with high WMH volumes and low SPPB scores, suggesting a possible mechanism of physical resilience within the brain.

## METHODS

### B-NET Study Design

Data are from the Brain Networks and Mobility (B-NET) study (NCT03430427). B-NET is an ongoing longitudinal, observational trial of community-dwelling older adults aged 70 and older recruited from Forsyth County, NC and surrounding regions. Participants were asked to complete a cognitive battery and various physical function tests at baseline with follow-up visits at 6, 18, and 30 months. Brain MRIs were collected at baseline and the 30-month follow-up visit. Baseline cross-sectional data are presented and 30-month collections are ongoing.

### Participants

In total, 192 participants were enrolled in B-NET. To be included, participants had to be above the age of 70, willing to sign informed consent, and able to communicate with study personnel. Exclusion criteria included being unwilling or unable to have a brain MRI, being a single or double amputee, having musculoskeletal implants severe enough to impede functional testing (e.g. joint replacements), or dependency on a walker or another person to ambulate. Participants were excluded for evidence of cognitive impairment determined by a combination of the Montreal Cognitive Assessment (MoCA) and additional cognitive testing as needed. Further exclusion criteria and a full description of cognitive inclusion criteria can be found here [20]. Three participants were excluded due to imaging artifacts resulting from excessive head motion; consequently, the resilience analyses presented here includes 189 participants. The study complied with the Declaration of Helsinki and written informed consent was obtained from each participant prior to any data collection. The Institutional Review Board (IRB) of the Wake Forest School of Medicine approved the study.

### Mobility Function Testing

Mobility function was tested using the expanded Short Physical Performance Battery (eSPPB) [21] and scores were calculated for both the eSPPB and SPPB [22]. The SPPB tests balance (stand with feet side-by-side, semi-tandem, and tandem for 10 seconds in each position), gait speed (4 meter walk at usual pace), and chair stand (time to rise from a chair 5 times). Each test is scored from 0-4, with 0 being the worst and 4 the best, for a total score range 0-12. The eSPPB increases the difficulty of the standing balance task by asking participants to hold postures for 30 seconds instead of 10 seconds and adds a one-leg stand. A narrow walk is also added to the testing in the eSPPB. Categorized scores of the SPPB were used to group participants into resilience groups (see below) given their associations with impairment [23]. In separate analyses, scores of the eSPPB were used for continuous analyses given its component scores are calculated based on the percentage of the best possible score (a continuous measure), not according to ranges of performance (a categorical measure). The resulting score is normally distributed, continuous and ranges from 0-4 rather than the traditional 12-point right-skewed categorical score distribution.

### Resilience Groupings

The groupings of WMH burden and SPPB scores were determined by a median split (Figure 1). For the SPPB, the median split fell at a score of 10, meaning that scores of 10 or higher (10-12) were considered “healthy” mobility while scores of 9 and below were considered “impaired” mobility. Participants in the top half of WMH burden (high burden) and bottom half of SPPB scores (impaired physical function) are termed Expected Impaired (EI) as they would be expected to have lower mobility function. Those in the bottom half of WMH burden (low burden) and top half of SPPB function (healthy physical function) are termed Expected Healthy (EH). Participants who were in the bottom half of WMH burden (low burden), yet also in the bottom half of SPPB scores (impaired physical function) are termed Unexpected Impaired (UI). Those in the top half of WMH burden who maintained high SPPB scores are termed Unexpected Healthy (UH). In the present study, we consider UH to show physical resilience as their physical function is healthy despite a large burden of white matter lesions. General group characteristics can be viewed in Table 1.

**Figure 1.**
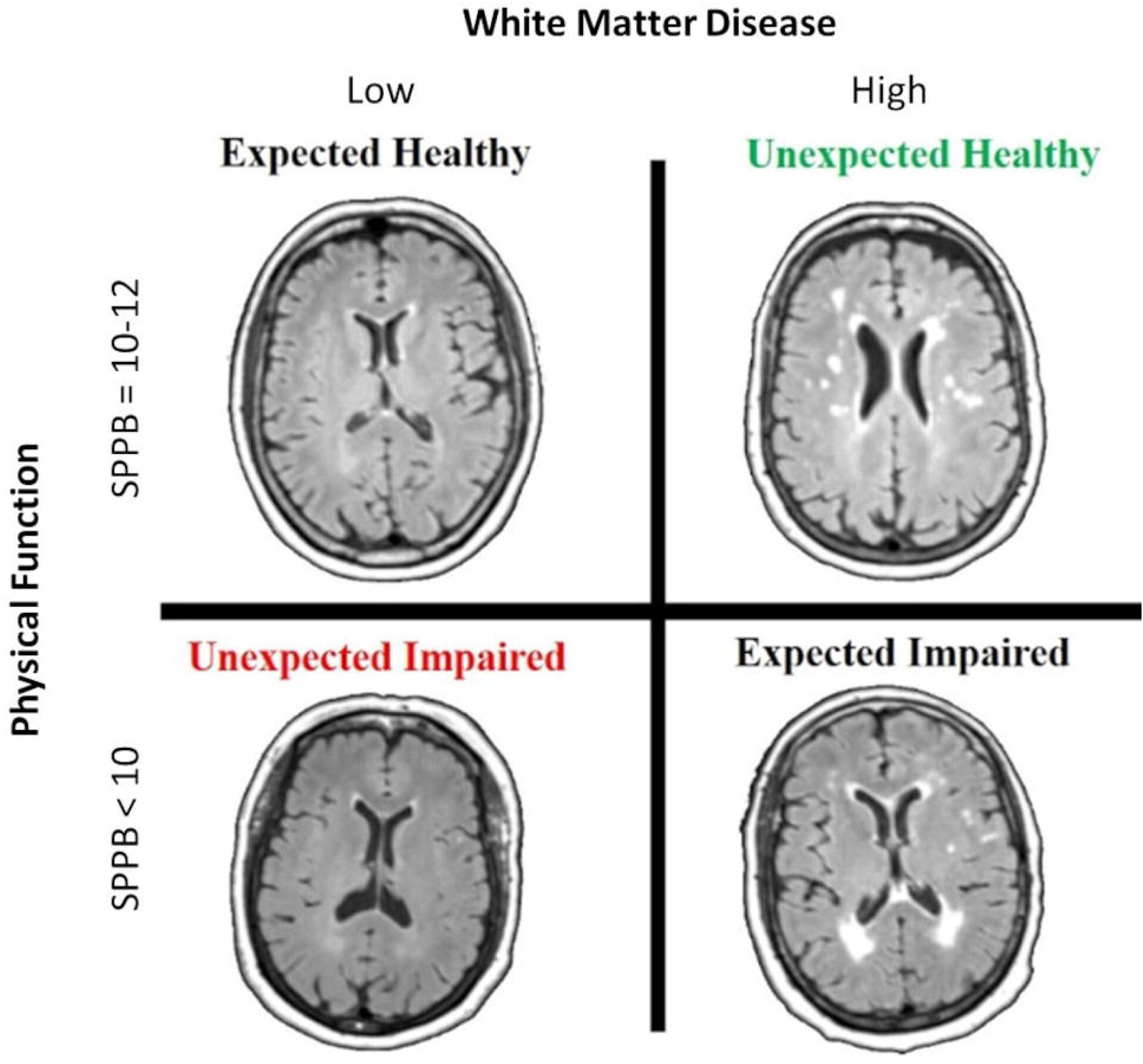
(Four Resilience Groups): Participants were grouped based on median splits of the short physical performance battery (SPPB) and white matter hyperintensity (WMH) volume. SPPB scores of 10-12 were considered “healthy mobility” while scores below 10 were considered “impaired mobility”. Expected Healthy (EH) had low WMH volumes and good mobility, Expected Impaired (EI) had high WMH volumes and poor mobility, Unexpected Impaired (UI) had low WMH volumes and poor mobility, and Unexpected Healthy (UH) had high WMH volumes and good mobility. UH are perceived to have high physical resilience.

**Table 1:** General Sample Characteristics.

|  | Full Sample<br>(N=189) | EH<br>(N=81) | EI<br>(N=42) | UI<br>(N=13) | UH<br>(N=53) | p-value |
| --- | --- | --- | --- | --- | --- | --- |
| Age (years ± STD) | 76.4 ± 4.7 | 74.5 ± 3.5 | 78.7 ± 4.8 | 77.4 ± 5.3 | 77.3 ± 4.9 | <0.01* |
| Education (years ± STD) | 15.7 ± 2.4 | 15.8 ± 2.3 | 15.4 ± 1.9 | 16.1 ± 3.5 | 15.5 ± 2.6 | 0.61 |
| Female N (%) | 105 (55.6) | 42 (51.9) | 23 (54.8) | 7 (53.9) | 32 (60.4) | 0.28 |
| White N (%) | 168 (88.9) | 74 (91.4) | 38 (90.5) | 10 (76.9) | 45 (84.9) | 0.56 |
| BMI (kg/m<sup>2</sup>) | 28.38 ± 5.64 | 27.81 ± 4.97 | 30.44 ± 6.16 | 32.45 ± 9.34 | 26.61 ± 3.65 | <0.01* |
| SPPB | 10.23 ± 1.60 | 10.99 ± 0.84 | 8.26 ± 0.87 | 7.77 ± 1.25 | 11.23 ± 0.82 | <0.01* |
| eSPPB | 2.01 ± 0.52 | 2.23 ± 0.46 | 1.54 ± 0.31 | 1.55 ± 0.55 | 2.17 ± 0.41 | <0.01* |
| 4-Meter GS (m/s) | 0.976 ± 0.200 | 1.065 ± 0.203 | 0.817 ± 0.130 | 0.777 ± 0.137 | 1.016 ± 0.136 | <0.01* |
| MoCA | 25.61 ± 2.20 | 25.81 ± 2.01 | 25.79 ± 2.34 | 24.77 ± 2.55 | 25.38 ± 2.21 | 0.42 |
| DSC | 55.15 ± 12.24 | 57.36 ± 12.20 | 51.19 ± 11.92 | 51.38 ± 9.51 | 55.83 ± 12.18 | 0.02* |
| Total WMH | 1.955 ± 0.997 | 1.134 ± 0.662 | 2.759 ± 0.753 | 1.607 ± 0.447 | 2.645 ± 0.596 | <0.01* |

\*significance at p<0.05
P-values from unadjusted ANOVAs, and chi-square tests for sex and race, estimating the group differences in general population characteristics
Note: EH=Expected Healthy, EI=Expected Impaired, UI=Unexpected Impaired, UH=Unexpected Healthy, BMI = Body Mass Index, SPPB = Short Physical Performance Battery, eSPPB = expanded Short Physical Performance Battery, GS = gait speed, MoCA = Montreal Cognitive Assessment, DSC = Digit Symbol Coding, WMH = log-normalized White Matter Hyperintensity volume

### Magnetic Resonance Imaging

All brain images were collected on a Siemens 3T Skyra MRI Scanner equipped with a 32-channel head coil. Each scan session lasted approximately one hour and includes structural and functional scans. The current study used the T1-weighted 3D volumetric MPRAGE anatomical image (TR=2300ms; TE=2.98ms; number of slices=192; voxel dimensions=1.0×1.0×1.0mm; FOV=256mm; scan duration=312s), T2 FLAIR images (TR=4800ms; TE=4.41ms; number of slices=160; voxel dimensions=1.0×1.0×1.2mm; FOV=256mm; scan duration=293s), and resting-state blood oxygenation level-dependent (BOLD) images (TR=2000ms; TE=25ms; number of slices=35; voxel dimensions=4.0×4.0×5.0mm; FOV=256mm; scan duration=460s). The BOLD images were collected parallel to the anterior commissure-posterior commissure line using multi-slice gradient-echo planar imaging [24].

### White Matter Hyperintensity Calculation

WMHs were identified on T2 FLAIR images using the lesion prediction algorithm (Schmidt, 2017, Chapter 6.1) implemented in the LST toolbox version 2.0.15 for Statistical Parametric Mapping version 12 (SPM12, http://www.fil.ion.ucl.ac.uk/spm). WMH were segmented by providing an estimate for the lesion probability for each voxel in the image. Next, the lesion probability maps were transformed into Montreal Neurologic Institute (MNI) brain space using Advanced Normalization Tools software (ANTs) [25] to account for variations in head size between participants. The total WMH volume for each participant was extracted using the fslstats command in FSL v5.0 (www.fmrib.ox.ac.uk/fsl) [26]. A log transform of the total WMH volume for each participants reduced the skewness of the data.

### fMRI Pre-processing

Structural image segmentation, including removal of non-brain tissue and cerebrospinal fluid (CSF), was completed using SPM12s described in [20]. Briefly, co-registration of the functional and anatomical images was improved through creation of a whole-brain mask for each participant using both white and grey matter segments. All images were manually inspected and cleaned to remove excess parenchymal tissue using MRIcron (https://www.nitrc.org/projects/mricron). High-resolution T1-weighted images were spatially normalized to the MNI template using ANTs [27]. Distortion correction was completed using FSL. The first 10 volumes were dropped to allow for signal stabilization. Slice time correction and realignment of functional images were completed using SPM12. BOLD images were co-registered to native-space anatomical images and warped to MNI space using the transformation derived from ANTs. Motion scrubbing was applied [28] and data were band-pass filtered (0.009 – 0.08 Hz) to account for low-frequency drift and physiologic noise. Confounding signals (white matter, grey matter, CSF, and 6 rigid-body motion parameters generated during the realignment process) were regressed out from the filtered data.

### Calculation of Brain Networks

The presence of a connection between two nodes, with each voxel representing a node in these analyses, (i and j) was determined by completing a time series regression analysis. This produced a cross-correlation matrix of Pearson’s correlation coefficients taken to represent the connectivity between every network node. A threshold was then chosen to dichotomize the data and create the final binary adjacency matrix (A_ij_) where values above the threshold indicate a connection is present (e.g. correlation coefficients are transformed to either a 0 indicating no connection or a 1 where there is a connection between nodes). A_ij_ is an n x n matrix where n is the number of brain voxels (∼20,000). This matrix notes the presence or absence of a connection between any two nodes (represented by i and j). The threshold, calculated as S = log (n)/log (K), is conserved to ensure comparable network density across participants. Here n represents the number of network nodes and the K is the average node degree, or the average number of edges connected to each node. Our networks were thresholded at S = 2.5 based on previous work showing that this threshold produces networks with density ratios that most closely compare to other naturally generated networks [29].

### Brain Network Analysis

Community structure analyses were performed on each participant’s brain network to assign each voxel to a network community. A community is defined as a group of nodes that have more connections with each other than to other nodes in the total network [17]. The strength of community structures was identified using network modularity or Q [30]. A dynamic Markov process [31] was used to maximize Q and partition brain networks into communities. As there is a stochastic component to the modularity algorithm, it was run 100 times and the partition associated with the highest Q value was chosen for that given study participant. Following partitioning of an individual’s brain network, each voxel was assigned the appropriate community membership and the spatial maps for the community organization were used in the second-level group analyses as described below.

Scaled Inclusivity (SI), a statistic utilized to compute the spatial consistency of community structure across a group of participants [32], was used for group analyses. Values of SI range between 0 and 1; perfect spatial alignment of a community across all participants would result in a value of 1 for all nodes in that community. In practice, communities are not made of exactly the same nodes across study participants as the spatial patterns vary across participants due to differences in the community organization. Thus, in practice each node is assigned a value less than 1 depending on the discordance across participants (for a detailed description of this method applied to the brain see [33]). High SI values indicate that a particular network community is stable across the group and occupies comparable brain regions in each person. SI values can be compared between groups or conditions to identify population differences in community organization.

Here, we performed SI analyses using regions-of-interest (ROIs) to define *a priori* intrinsic brain subnetworks as described in [20]. In the current study, the SMN was chosen as the primary ROI due to its established involvement in motor function and our prior work establishing a relationship between higher scores of the SPPB and a more consistent SMN-CS [17]. SI was computed across all participants. SI analyses were performed to compare SMN-CS between reserve groups. Analyses were also performed to identify second-order connectivity to determine how the sensorimotor cortex links to other brain regions. A first-order connection of node i in sensorimotor cortex is any node j that has a direct connection to node i. A second-order connection of node i is any node (k) that is directly connected to a first-order connection of node i (i.e. node j); note that node k ≠ i. For more details, see [17]. A log-transform of both community structure and second-order connections for each participant reduced the skewedness of the data.

### Statistical Analyses

The effect of resilience group on functional network properties was explored using a cell means parameterization of a generalized linear model with age, sex, and race as covariates and using contrasts to test main effects, interactions and perform pairwise testing between groups (Proc GLM, SAS version 9.4, SAS Institute Inc., Cary, NC). Pairwise group differences between adjusted means (and 95% Confidence Intervals) in network measures were estimated with the Tukey-Kramer procedure used to correct for multiple comparisons. Pairwise differences in the 3-dimensional spatial distribution of second-order connections by groups was assessed using a permutation statistic [34].

In order to better understand the relationships between functional brain network measures and physical and cognitive function, a separate set of linear models including all 189 participants was used to estimate the associations between continuous network variables (dependent variables), and physical function, and cognition as predictors. Physical function was represented using the continuous measures for eSPPB and gait speed rather than using median splits. Cognition was represented using scores on the MoCA, and digit symbol coding task (DSC). Model 1 adjusted for age, self-reported sex, self-reported race, and body mass index (BMI) as covariates while Model 2 added WMH volume as a covariate. Sensitivity analyses were conducted with both network measures (community structure and second-order connections) to assess the impact of influential outliers and confirm model robustness.

## RESULTS

### Participant Characteristics

Of the 189 participants (mean age=76.5±4.7; female=52.4%, Caucasian=78.9%; mean years of education=15.8±2.4), 81 were characterized as Expected Healthy (EH), 42 as Expected Impaired (EI), 13 as Unexpected Impaired (UI), and 53 as Unexpected Healthy (UH). A detailed description of participant characteristics can be found in Table 1. Adults with better SPPB scores (EH and UH) were, on average, younger (p<0.01), had lower BMI (p<0.01), and scored higher on the DSC (p=0.02) than those with low SPPB scores (EI and UI). No significant between-group differences were observed in education, sex, race, or MoCA scores.

### Community Structure

Analyses of the effect of resilience grouping on community structure identified a significant difference in means (p=0.013), and using Tukey-Kramer adjustments, we were able to conclude that the EH and UI groups were different (see Figure 3 for pairwise differences adjusted for age, sex and race). Contrasts applied to these means identified main effects for SPPB (p=0.003), no main effect for WMH group (p=0.80) and no interaction effect (p=0.14). Group differences in SMN-CS are visualized using brain maps in Figure 2.

**Figure 2.**
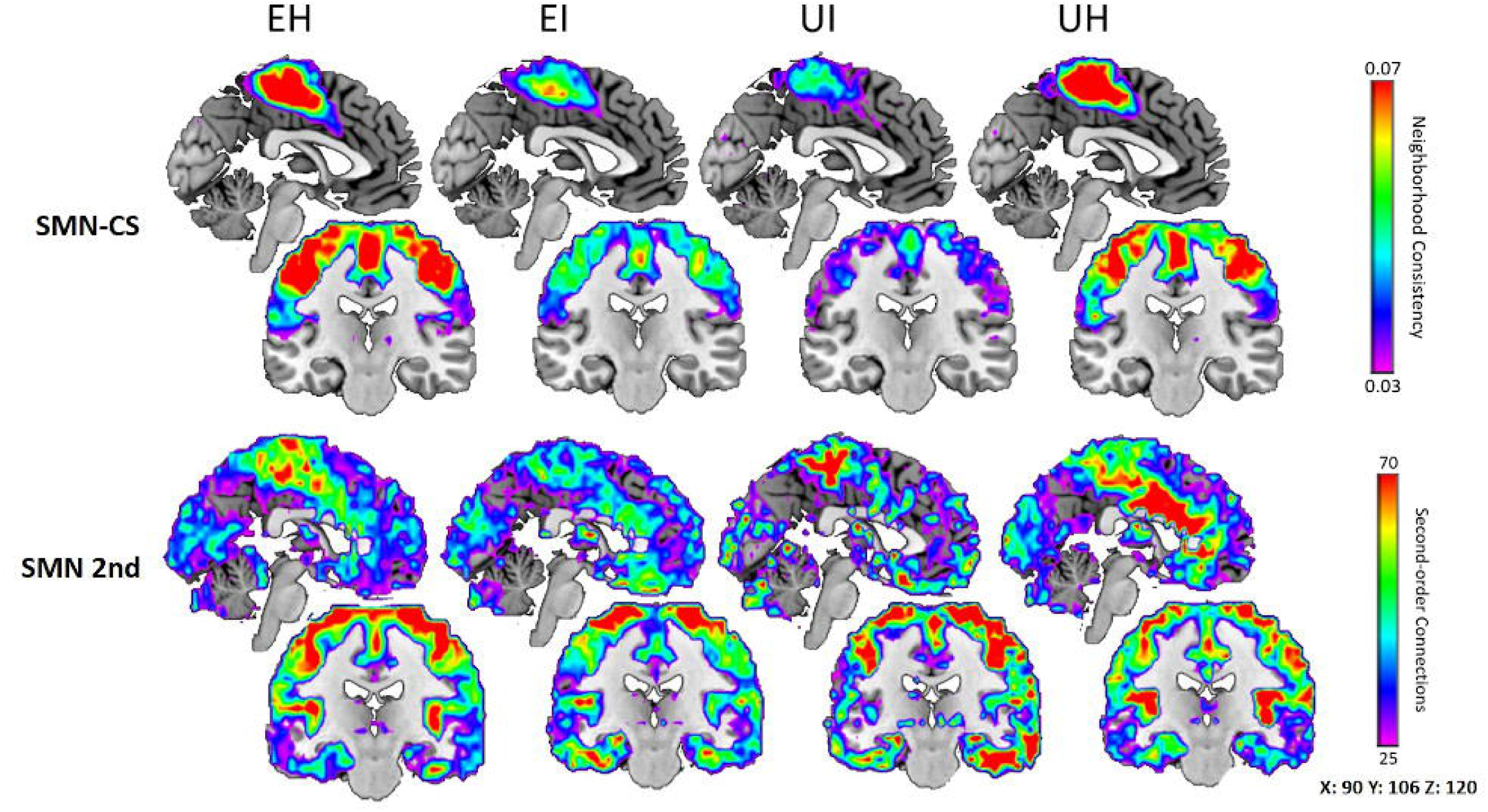
(Resilience Group Network Comparison): Brain maps showing the resting-state consistency of the sensorimotor cortex (SMN-CS, top row) and second-order connections from the SMN to the rest of the network (SMN 2nd, bottom row) between Expected Healthy (EH), Expected Impaired (EI), Unexpected Impaired (UI), and Unexpected Healthy (UH). For SMN-CS, cool colors indicate a less consistent network community structure than warm colors. At rest, EH and UH had a more consistent SMN-CS than EI and UI. For SMN 2nd, cool colors indicate fewer second-order connections coming from the somatomotor cortex (SMN) than warm colors. UH showed a unique pattern of connectivity in that they had higher numbers of second-order connections from the SMN to the anterior cingulate cortex (ACC) than all other groups, particularly EI.

**Figure 3.**
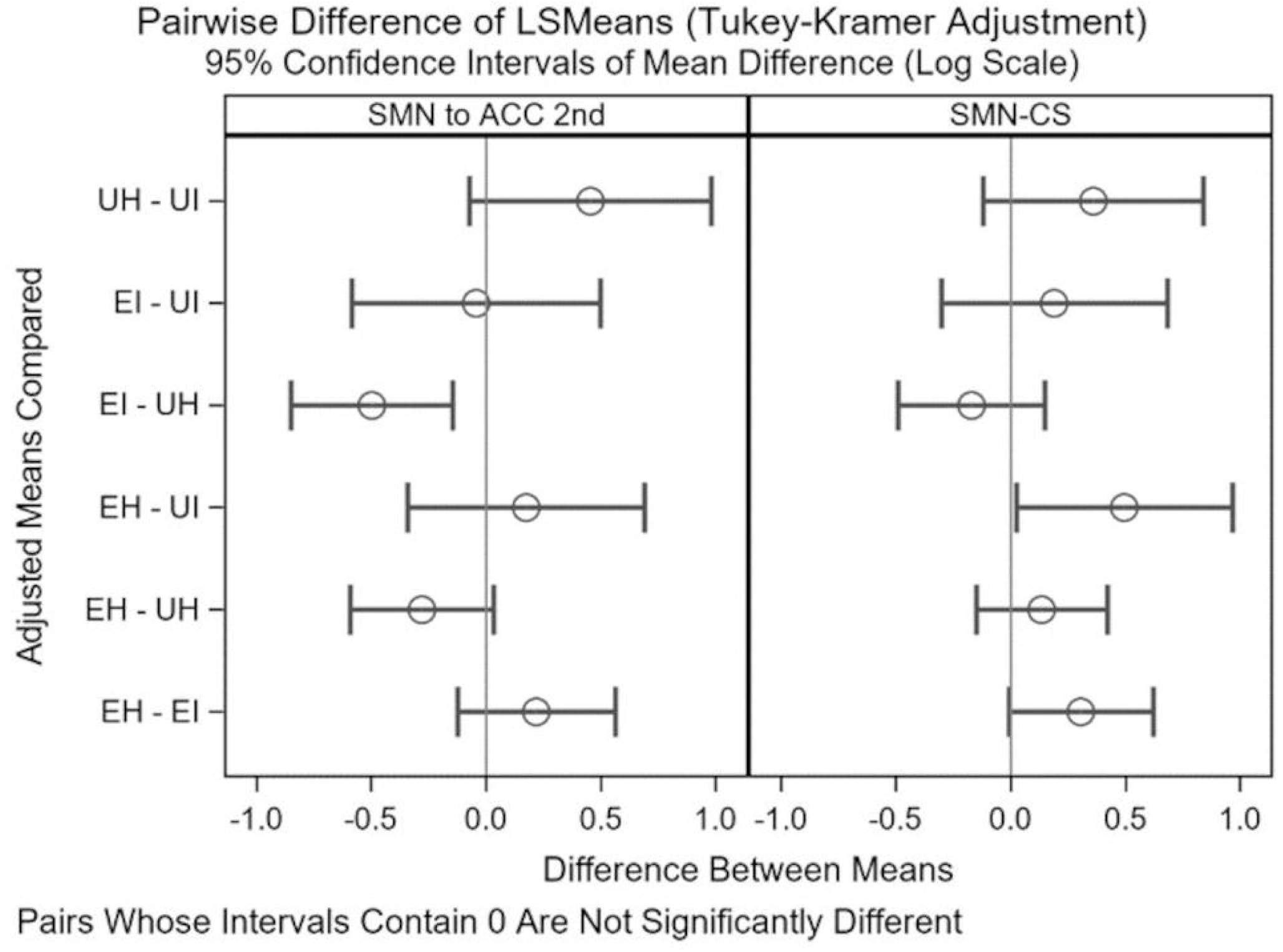
(Pairwise Differences in Least Square Means (LSMeans): Output from a cell means parameterization of a generalized linear model with age, sex, and race as covariates and using contrasts to test main effects, interactions and perform pairwise testing between groups. A significant difference in SMN-CS means, using Tukey-Kramer adjustments, was identified such that EH and UI were different. A significant difference in second-order connection means was reported such that EI and UH were different. For both network variables, a main effect was identified for SPPB, but not for WMH group or the interaction effect.

### Second-order Connections

We next evaluated the patterns of connectivity from SMN to the rest of the brain adjusting for age, sex, and race (see Figure 2 for network community structure and second-order connection maps, and Figure 3 for pairwise differences adjusted for age, sex and race). A significant (p<0.05) pairwise spatial difference in second-order connectivity was identified with the permutation statistic such that UH showed a pattern of second-order connections from SMN that was distinct from EI. Although the permutation test compares the whole-brain connectivity patterns, it is evident from the results that 2^nd^ order connectivity to the ACC is particularly high in the UH group. Analyses of the effect of resilience grouping on second-order connections from SMN to ACC identified an overall significant difference in means (p=0.002), and using Tukey-Kramer adjustments, we were able to conclude that the EI and UH groups were different (See Figure 3). Contrasts applied to these means identified main effects for SPPB (p=0.006), no main effect for WMH group (p=0.33) and no interaction effect (p=0.18)

### Associations with Physical and Cognitive Function

Table 2 contains the output of the linear models exploring associations between continuous network measures, physical function, and cognition. SMN-CS was not significantly associated with eSPPB scores (Model 1: p=0.085, Model 2: p=0.129), but did show a significant positive association with gait speed (Model 1: p<0.01, Model 2: p=0.019). Higher numbers of SMN second-order connections to the ACC were observed with higher eSPPB scores (Model 1: p=0.106, Model 2: p=0.060) and faster gait speed (Model 1: p=0.207, Model 2: p=0.117), but these associations did not reach significance. MoCA scores showed no significant associations with SMN-CS (Model 1: p=0.705, Model 2: p=0.836) or SMN second-order connections to the ACC (Model 1: p=0.454, Model 2: p=0.494). DSC scores were not associated with SMN-CS (Model 1: p=0.068, Model 2: p=0.105). However, an inverse association between DSC scores and the number of second-order connections from SMN to ACC was found (Model 1: p=0.022, Model 2: p=0.034), but the effect size was small.

**Table 2:** Output From Linear Models Exploring Associations Between Continuous Functional Network Measures and Scores of Physical and Cognitive Function.

| | Model | SMN to ACC ($\beta$, 95% CI, T-stat, p-value) | SMN-CS ($\beta$, 95% CI, T-stat, p-value) |
| --- | --- | --- | --- |
| DSC | 1 | -0.010 (-0.018, -0.001); -2.3; 0.022 | 0.007 (-0.001, 0.014); 1.84; 0.068 |
|  | 2 | -0.009 (-0.017, -0.001); -2.1; 0.034 | 0.006 (-0.001, 0.014); 1.63; 0.105 |
| eSPPB | 1 | 0.177 (-0.038, 0.391); 1.62; 0.106 | 0.170 (-0.023, 0.363); 1.73; 0.085 |
|  | 2 | 0.207 (-0.009, 0.424); 1.89; 0.060 | 0.152 (-0.045, 0.349); 1.52; 0.129 |
| Gait Speed (m/s) | 1 | 0.351 (-0.196, 0.898); 1.27; 0.207 | 0.651 (0.167, 1.136); 2.65; 0.009 |
|  | 2 | 0.444 (-0.112, 1.000); 1.57; 0.117 | 0.598 (0.098, 1.098); 2.36; 0.019 |
| MOCA | 1 | 0.017 (-0.028, 0.062); 0.75; 0.454 | 0.008 (-0.033, 0.048); 0.38; 0.705 |
|  | 2 | 0.016 (-0.030, 0.061); 0.68; 0.494 | 0.004 (-0.037, 0.045); 0.21; 0.836 |

\*significance at $p < 0.05$
Model 1 (Physical Function and Cognition): Effect of network measures on physical and cognitive function adjusted for age, sex, race, and BMI
Model 2 (Physical Function and Cognition): Model 1 + WMH volume
Note: SPPB = Short Physical Performance Battery, eSPPB = expanded SPPB, MoCA = Montreal Cognitive Assessment, DSC = Digit Symbol Coding Test, SMN to ACC = Second-order connections from Sensorimotor Cortex to Anterior Cingulate Cortex, SMN-CS = Sensorimotor Cortex Community Structure

## DISCUSSION

In the present study, we investigated the neural correlates of physical resilience defined as having good mobility function in the face of a higher WMH burden. We used graph theory network analyses grouping participants based on WMH volumes and SPPB scores. We hypothesized that individuals with high brain lesion load but healthy mobility (high resilience - UH) would show compensatory shifts in functional connectivity when compared with their non-resilient counterparts (EI). We also hypothesized that these shifts would provide indications of physical resilience within the brain. Permutation analyses on whole brain images indicated that UH, characterized as having high physical resilience, displayed a unique pattern of second-order connections from the SMN to the ACC when compared with EI. Pairwise comparisons of SMN-CS showed that UH was less consistent than EH, but more consistent than both EI and UI. UH also displayed higher numbers of second-order connections from SMN to ACC than all other groups, and this was most clear cut between UH and EI. This pattern of connectivity in UH, who maintained high levels of mobility despite high WMH burdens, is possibly a mechanism underlying physical resilience within the brain. Although no significant effect of the resilience interaction term was found in the regression analyses, the main effects indicated that higher SPPB scores are associated with increased SMN-CS and second-order connections from SMN to ACC. We did not observe significant relationships between network measures and scores of cognitive and physical function in our association analyses, but this may be due to our relatively low sample size in some groups.

The high number of second-order connections from SMN to ACC in UH compared to EI appears to confirm our hypothesis that resilient individuals would display compensatory shifts in functional connectivity when compared to their non-resilient counterparts. Functionally, the ACC plays primary roles in attentional allocation and control as well as intentional motor control [35] and decision-making [34]. Structurally, the ACC is highly integrated with cortical motor regions [36]. Recently, the ACC has gained attention as a key region in a “fine-tuning” network for human locomotion where continuous flexibility and adaptability is necessary [19]. The ACC is also a key node in a prominent brain network, the salience network (SAL) [34]. Generally, the SAL plays major roles in the initiation of cognitive control [35], organization of behavioral responses [36], and detection and integration of emotional and sensory stimuli [37]. The SAL’s role in cognitive control may be particularly important for resilience as it centers around its capacity to drive the alternation between DMN (internally directed cognition) and central executive network (externally directed cognition), which allows for more efficient navigation through one’s environment.

Interestingly, the SAL is the sole host to a unique class of recently discovered neurons: von Economo neurons (VENs). While VENs are found in both the ACC and fronto-insular cortex, they are more densely populated in the ACC. VENs are thought to provide the rapid relay of neuronal signals to other brain regions and thus provide the basis for cognitive control [38, 39]. While our understanding of VENs is limited, they appear to be unique to humans and apes and emerge mainly after birth until the age of approximately 4 years [38, 39]. Importantly, higher volumes and densities of VENs are associated with resilience and reduced vulnerability to Alzheimer’s pathology [40]. In addition to further highlighting the potential for second-order connections between SMN and SAL to be a resilient mechanism, these findings also suggest that this pattern of connectivity, possibly related to VENs, may be apparent throughout the lifespan.

Altogether, we observed that UH, who are perceived as having high physical resilience, displayed a unique pattern of connectivity. This suggests that this ability to form higher numbers of connections despite increased pathologic burden, a marker for neural plasticity, could be a compensatory mechanism for the maintenance of physical function. More specifically, given the role of the ACC and SAL in motor and cognitive control as well as stimuli integration, increased connectivity from the SMN to these regions could be a marker of CNS resilience, which is one component of physical resilience.

Community structure analyses did not reveal any significant differences in SMN-CS of UH to any of the other three groups. In fact, the only significant group differences observed were EH being more consistent than UI, which is in line with our previous findings on SMN-CS and SPPB scores [17]. While not significant, SMN-CS consistency of UH was most similar to that of EH in that the average consistency was higher than that of both impaired groups, which further highlights the positive relationship between SMN-CS and SPPB scores. The lack of impairment in SMN connectivity in UH could indicate that they are resistant to the deleterious effects of WMH within the SMN. An important and unanswered question, however, is if maintenance of SMN-CS alone is enough or whether resilient individuals also need to maintain the increased level of second-order connections to the ACC to maintain mobility. Of note, UH also did not show any significant cognitive differences from EH despite the increased burden of WMH.

While imaging findings are compelling, mobility requires energetic input, structural integrity, muscle strength, and good sensory function among other factors. Therefore, physical resilience is likely comprised of multiple reserves or resiliencies [1]. For example, an individual might have reserve within a given system if they have a high level of mitochondrial health due to good health habits or genetic variation that acts as a buffer against age-related declines in mitochondrial function and prolonging mobility. Alternately, reserve in one system might compensate for or buffer against loss in another. For example, bone structural integrity is weakened by joint damage that may require a joint replacement, but good muscle strength from lifelong exercise habits helps maintain and recover mobility more rapidly.

Our study benefits from several key strengths. First, the study design of B-NET provides advantages in that it consists of a large, well-characterized cohort of community-dwelling older adults who are free of mild cognitive impairment and known central nervous disease. The wealth of testing data provides unique opportunities to advance our understanding of mobility-related functional brain networks. Our study was also strengthened by the use of WMH and SPPB scores to characterize physical resilience in combination with functional network architecture. To our knowledge, we are the first to make use of this paradigm, which may identify individuals with high resilience.

There were also several limitations in the current study. First, the data presented here are cross-sectional, which limits our ability to comment on longitudinal effects. However, 30-month follow-up MRI and fMRI scans will be collected and allow for future longitudinal network analyses. In addition, our sample was recruited as cognitively normal. While this does provide opportunities to explore associations in the absence of cognitive impairment, we are not able to compare network connectivity between individuals who are cognitively normal vs. impaired. Finally, we were limited statistically by the reduced sample size within our resilience groups because of our resilience grouping method. While our total sample size was 189, within group sizes ranged from 81 to 13, which limited statistical power despite the visually apparent group differences in network measures. It is possible that our analyses would have revealed stronger associations between the resilience interaction term and network measures with larger within group samples. Future analyses should aim to recruit larger, equally distributed samples to address this limitation.

## CONCLUSION

Utilizing graph theory to study brain network architecture in a large, well-characterized cohort of community-dwelling older adults, we have identified a shift in functional connectivity of resilient participants that may be a marker of physical resilience in the brain. Increased connectivity from SMN to ACC appears to be a driving factor for resilient individuals to maintain physical function and may be a key node of a “resilience brain network”. Additionally, our paradigm of combining WMH volume with scores of the SPPB is easily replicable and proved effective in identifying individuals with high physical resilience. Taken with the previous literature showing that neural resources are shared between mobility and cognition [6, 41], the property of resilience may not be specific only to physical function but shared with cognition. Future studies should aim to replicate and expand on these analyses in populations that are cognitively normal as well as impaired, by studying different brain pathologies, and by studying the differences in group responses to interventions.

## ACKNOWLEDGMENTS

The authors would like to thank study coordinators and MRI staff for their efforts in collecting data. We would also like to thank study participants for taking part in this research.

## AUTHOR CONTRIBUTIONS

Conceptualization, PJL, SBK, and CEH; Methodology, PJL, SBK, MEM, SNL, CEH, RGL, and BRN; Validation, PJL, SBK, MEM, SNL, CEH, RGL, and BRN; Investigation, PJL, SBK, SNL, MEM, CEH, LDB, and BRN; Writing: BRN, CEH, SNL, PJL, SKB, MEM, EPH, and LDB; Funding Acquisition, SKB and PJL. All co-authors read and agreed to the published version of the manuscript.

## CONFLICTS OF INTEREST

The authors declare no conflicts of interest.

## Notes

### Competing Interest Statement

The authors have declared no competing interest.

